# Faster is better: Visual responses to motion are stronger for higher refresh rates

**DOI:** 10.1101/505354

**Authors:** Mina A. Khoei, Francesco Galluppi, Quentin Sabatier, Pierre Pouget, Benoit R Cottereau, Ryad Benosman

## Abstract

Although neural responses with a millisecond precision were reported in the retina, lateral geniculate nucleus and visual cortex of multiple species, the presence and role of such a fine temporal structure is still debated at the cortical level and the general belief remains that early visual system encodes information at slower timescales. In this study, we used a new stimulation platform to generate visual stimuli that were very precisely encoded in time and we characterized in human subjects the EEG responses to moving patterns that shared the same global motion but differed in their fine scale spatio-temporal properties. In two experiments, we manipulated the information within temporal windows that corresponded to the frame duration in conventional (1/60 = 16.7ms, experience 1) and more recent (1/120 = 8.3ms, experience 2) visual displays. Our results demonstrate that EEG responses to temporally dense and coherent trajectories are significantly stronger than those to control conditions without these properties. They extend previous results from our group that showed that accurate temporal information (<10ms) significantly improve perceptual judgments on spatial discrimination, digit recognition and sensitivity for speed [Kime et al., 2016]. Altogether, our results suggest that instead of low-pass filtering the temporal information it receives from its thalamic afferents, the visual cortex may actually exploit its richness to improve visual perception.

**Significance statement:** Most visual experiments use frame-based stimulation screens which are synchronous and slow. This might not engage the full encoding capacity of the visual system, as in natural condition, visual information is asynchronously acquired by the retina with a very high temporal resolution. In the present study, we used an unconventional and highly configurable stimulation platform to determine the EEG responses to moving patterns that shared the same global motion but differed in their fine scale (<10ms) spatio-temporal properties. Our results demonstrate that brain responses in human are significantly stronger for motion patterns that are refreshed at very high frame rate (360Hz) and thereby provide a strong support for the use of very precise temporal stimulations in vision studies.

## 1 Introduction

Brain science has been slowed down in its quest to understand functionalities of the visual system by technical limitations of current display technologies that set refresh rates around the historical flicker-fusion of 24 frames per second. Current monitors and televisions have a maximum refresh rate in the range of 60-120Hz to match existing bottlenecks of broadcasting bandwidth and storage capacities. Although 24 fps is enough for most of human observers to have a stable perception of dynamic scene, it has been reported that such low frame rates induce dizziness and motion sickness when dealing with dynamic inputs. In addition, response of neurons in the early visual cortex of humans and primates have been reported to be aligned to the refresh rates of displayed video [Williams et al., 2004] at rates below 100Hz. LGN and V1 cells of cat also have demonstrated phase locked firing to refresh rate of conventional display monitors [Wollman and Palmer, 1995].

There has been little research on understanding the responsiveness of the visual system to higher frame rates and finer temporal resolutions and possible perceptive enhancements in experimental platforms. In most visual experiments, stimuli are displayed at middle or low frame rates, rarely higher than 200Hz. Evidently, daily human tasks go far beyond watching TV and delicate and demanding actions like driving require much higher spatio-temporal precision. Some studies suggest that visual system uses precise timed signaling from the retina, to the lateral geniculate nucleus (LGN) [Reinagel and Reid, 2002] and to the visual cortex, [Butts et al., 2007]. At the retina level, precise temporal response is known to be a native property of retina ganglion cells (RGC), holding over a wide range of stimulus patterns and contrast in different species [Gollisch and Meister, 2010, Berry et al., 1997]. Similarly, stimulus dependent temporal precision has been observed by in vivo recordings on cat LGN in response to drifting sinusoidal gratings [Reich et al., 1997]. In macaque, spiking patterns within higher visual area MT are clearly different for rigidly translating stimuli and highly dynamic random motion, showing a stimulus locked temporal modulation that disappears as the coherence level of motion in random dot patterns increases [Bair and Koch, 1996].

Some of these studies used subjective psychophysical reports in humans to determine if there is a frame rate threshold above which human perception does not improve anymore [Kuroki et al., 2007]. Others were based on interactive scenes and games to evaluate player performance as a function of frame rate [Claypool et al., 2006]. Recently, our group reported significant improvement of perceptual judgments on spatial discrimination, digit recognition and sensitivity to speed by increasing frame rates from 60Hz to 1KHz [Kime et al., 2016].

Existing studies share the general point of view that the refresh rate is encoded as an artefact on neural responses, without initiating any cortex-based mechanism. If millisecond temporal structure is important for the representation of visual information, one might expect it to be preserved and observable at the cortical level. In this study we used high temporal precisely controlled moving stimuli (random dot patterns) using a display system able to cover a wide range of stimuli from 30-1KHz. We observe that a high temporal precision stimulus influences EEG brain responses to global motion. This reveals that a high precision temporal structure is important for the representation of dynamic visual information in the human brain.

## 2 Material and Methods

The following section describes the methodology used to design and perform the experiments.

### 2.1 Subjects

Nineteen subjects (14 men, 5 women, age range 22-35 years old) participated in both experiments and had normal or corrected to normal vision. They were given detailed instruction about the experiment. This study was approved by the local ethic committee (IRB number 20122800001072) based notably on its compliance with the Helsinki Declaration.

### 2.2 Stimulation platform

Our stimuli were displayed using an evolution of the platform described in [Kime et al., 2016] to deliver precise-time controlled light stimulation at very high frame rates. The projecting system consisted in a Texas Instrument LightCrafter projector that controlled a DLP3000 Digital Micromirror Device (DMD). The DMD comprised an array of mirrors that could rapidly (1440 Hz, 0.7ms) switch between two discrete angular positions (−12° and 12°) to enable or disable the reflection of a light source on a screen [Sampsell, 1994]. The DLP3000 was composed of an array of 608 × 684 mirrors tilted by 45 degrees. We addressed every other line in the mirror array so as to avoid spatial artifacts which might be introduced by the native addressing scheme of the DMD. The DMD was controlled with an embedded Linux system preparing the bitplanes. The device was driven in binary mode, so as to independently control the on and off positions of the mirrors with a minimum time step of 0.7ms. Light intensity was therefore encoded with two binary values, corresponding to a mirror being on or off. The DMD platform projected the visual stimuli on a screen localized at 140cm in front of the participants. The projection covered a surface of 40 x 32cm (i.e. 16.3° × 13° of visual angle) and used a QVGA resolution of 340 × 240 pixels (i.e. of 1.3 × 1.3*mm* which corresponds to 0.05° of visual angle). Stimuli (40cd/m2) were projected in a dark room, with a light source of 530nm wavelengths against a dark background (9 cd/m2), leading to a Michelson contrast of 63.27%.

### 2.3 Stimulus

The two experiments of this study are based on frequency-encoded stimuli that contain the same global motion pattern but differ in their fine-scale spatio-temporal properties. For the first experiment, the three conditions were all derived from a stimulus that is classically used to characterize motion selective responses in primate visual cortex (see e.g. [Newsome and Pare, 1988]). This stimulus consisted in a circular 16.3 *degree*^2^ pattern of random dots (dot size: 0.05°, density: 1.4 dots/*degree*^2^) whose motion alternated at F = 3Hz between a fully coherent state and an incoherent state. During the coherent state (i.e. 166.7ms), all the dots moved in the same direction at a constant speed of 19.6°/*s*. The direction alternated between left (on odd coherent states) and right (on even coherent states). During the incoherent state, each dot moved along a direction drawn from a uniform distribution. Our three conditions consisted in different versions of this coherent/incoherent motion stimulus specifically designed to evaluate the impact of temporal precision on the brain responses recorded with EEG. For these three conditions, each dot had a displacement of 6 pixels during a 60Hz frame period (i.e. during 24*T* ≈ 16.6ms). In our main condition (360Hz), the dot positions were refreshed each time they moved by one pixel. Given our experimental setup (i.e. the projection resolution and dot speed), it corresponds to a refresh rate of 360Hz (i.e. dots were refreshed every 4Δ*T*). In the 60Hz condition, the dot position was updated at 60Hz (i.e. every 24Δ*T*)) and remained constant throughout the whole 60Hz frame period. In the random (RND-60) condition, the dot position was also updated every 4Δ*T* but the spatial positions of each dot along its trajectory were scrambled within each 60Hz frame period. These three conditions are illustrated on figure 1. Note that they all contain the same global motion pattern (i.e. the F = 3Hz transition between the coherent and incoherent motion states). The 60Hz and 360Hz conditions only differed in their refresh rate. If 60Hz monitors generate visual responses that are equivalent to those that are elicited in every day life (see the introduction), these two conditions should generate identical EEG responses. The 360Hz and RND-60 condition only differed in their spatio-temporal properties within a 60Hz frame period but had otherwise identical properties (i.e. refresh rate and spatial position of the dots during a complete state). If the retino-geniculate pathway filters the visual inputs to uniquely transmit the lowest temporal frequencies (i.e. < 100Hz) to the cortex (see the introduction), these two conditions should generate identical EEG responses. For this first experiment, stimuli were presented during trials of 15s and separated by 5s of inter-stimulus interval (ISI). For a given trial, the stimulus was randomly picked between the 3 experimental conditions (i.e. 360Hz, 60Hz and RND-60). Trials were grouped within recording block of 13 minutes and each block contained 39 trials in total. The entire recording session consisted in 3 blocks (117 trials in total, 39 for each condition) and approximately last 40 minutes. To further characterize the influence of fine-scale spatio-temporal properties on global motion responses, we recorded the responses to a second set of stimuli that were identical to those of the first experiment except that the two controls were derived from a refresh rate of 120Hz. These two controls conditions are therefore labeled as 120Hz and RND-120. Apart from this change in the two controls, this second experiment had the same properties (number of blocks, trials see above) as the first experiment.

**Figure 1:**
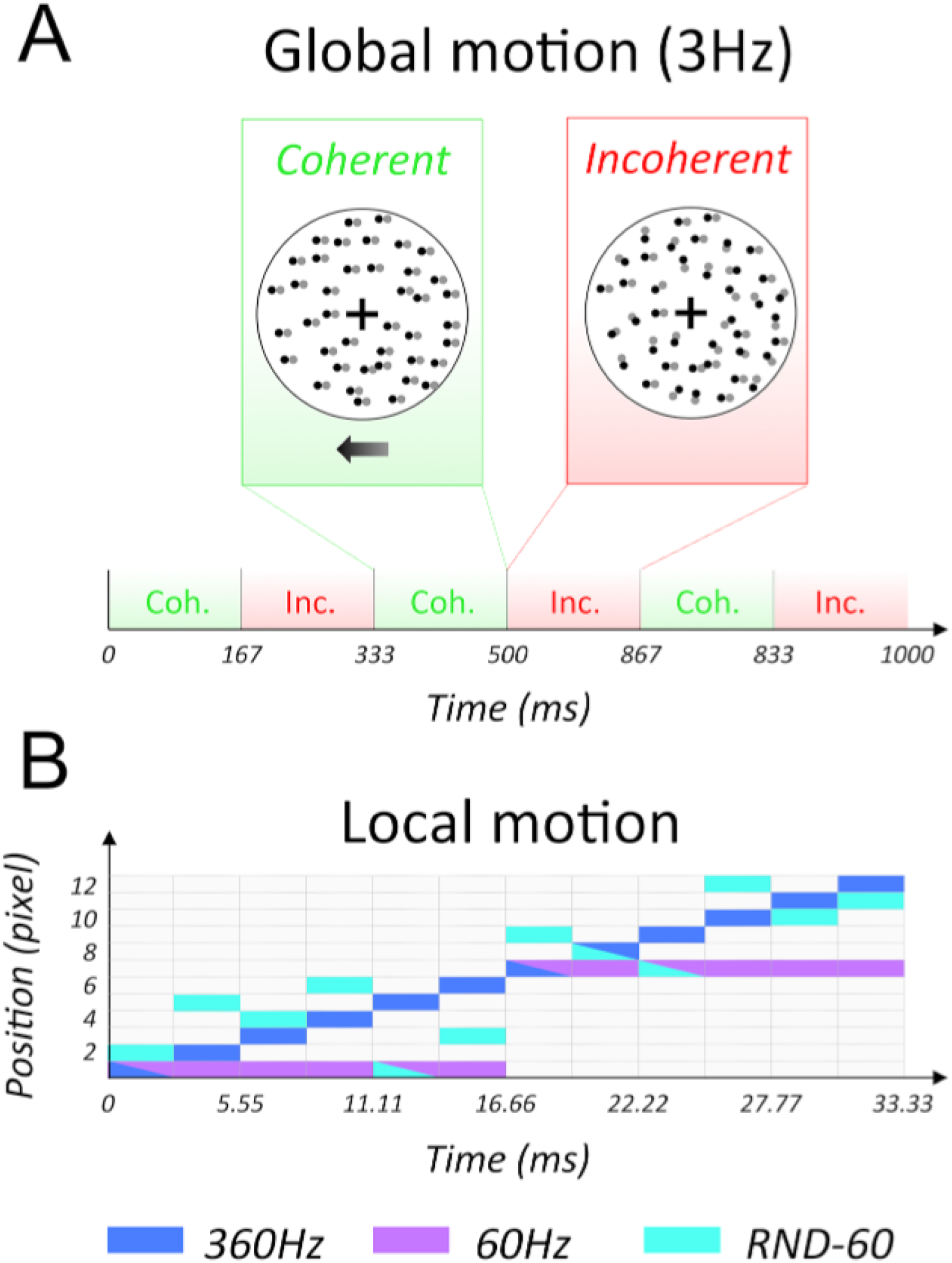
Properties of the three experimental conditions used in the first experiment. A) General temporal structure of the stimuli. The global motion of the dots alternates between a coherent state (in green) and an incoherent state (in red) at F = 3Hz. During the coherent states, all the dots move either leftward (on odd coherent states) or rightward (on even coherent states). During the incoherent state, each dot moves along a random direction. The color of the dots indicates their position at time t (in grey) and *t* + Δ*T* (in black). B) At a finer temporal scale, dot positions in the 360Hz condition are refreshed every 4Δ*T* (blue) whereas they are refreshed every 24Δ*T* in the 60Hz control condition (purple). In the RND-60 condition (cyan) dot positions are refreshed every 4Δ*T* but are scrambled within time windows of 24Δ*T*. All three conditions have the same global motion patterns at 3Hz but they differ in their fine-scale spatiotemporal properties.

### 2.4 EEG signal acquisition and pre-processing

EEG data were collected with a 64-sensor Biosemi system and processed with the Eeglab toolbox [Delorme and Makeig, 2004]. Raw data were first band-pass filtered offline, between 0.1 to 200 Hz, and then analysed to detect and exclude noisy or artifacted electrodes using eeglab probability methods and visual inspection. These electrodes were substituted by the average of their six nearest neighbours. On average, less than 10% of the electrodes were substituted; these electrodes were mainly located near the forehead or the ears. As we shall see below, in the context of our visual experiment, our stimuli mostly elicited activations in occipital and parietal electrodes. Therefore, this substitution approach had a negligible impact on our results. After this step, the EEG data was re-referenced to the common average of all 64 electrodes and divided into 3 datasets, one for each experimental condition. Each dataset was segmented into 1s long epochs, each epoch containing exactly 3 cycles of coherent versus incoherent motions. The first two epochs of each trial were discarded so as to eliminate the transient visual responses that are elicited by the onset of visual stimulation. EEG epochs that contained a large percentage of data samples exceeding a noise threshold (depending on the subject and ranging between 60 and 80 *μV*) were excluded from the analysis on a sensor-by-sensor basis. This was typically the case for epochs containing artifacts, such as blinks or eye movements. In addition, we also applied a probability based epoch rejection method that used the joint probability of the measured activities at each time point. This approach uses the fact that observing large absolute values at most electrodes is highly improbable and thereby probably reflects the presence of an artefact. In this case, our electrode threshold was the joint probability activity limit (in std. dev.) for an electrode.

The use of steady-state stimulation drives cortical responses at specific frequencies directly tied to the stimulus frequency [Norcia et al., 2015]. It is thus appropriate to quantify these responses in the frequency domain. Therefore, a Fourier analysis was applied on every remaining epoch using a discrete Fourier transform with a rectangular window. Given the time length of an epoch (i.e., 1s), this Fourier transformation led to a frequency resolution of 1Hz. For each frequency bin, the Fourier coefficients were then averaged across all the epochs and all the trials. Thus these average Fourier coefficients for a given condition were obtained from up to 507 values (i.e. 39 trials by condition, each 13 epochs, as the two first epochs of each trials were discarded, see above). For some subjects (5 in the first experiment, 1 in the second experiment), our pre-processing pipeline led to a limited average number of correct epochs (i.e. n < 150) for one or several conditions. Because in this case, Fourier coefficient estimation can be inaccurate, we discarded those subjects from further analyses. Our results are therefore based on n = 14 subjects for the first experiment and n = 18 subjects for the second one.

### 2.5 Signal-to-noise ratio analysis

We were particularly interested in the response magnitudes (i.e., the modules of the Fourier coefficients) at the odd (first and third) and even (second and fourth) components of the global motion frequency (1F = 3Hz and 3F = 9Hz for odd components and 2F = 6Hz and 4F = 12Hz for even components). This choice was motivated by the fact that significant responses were still observable in our spectra for the third and fourth harmonics (see e.g. figures 2 and 3). Both the first and third components reflect the asymmetries in the neural responses to our stimuli, i.e. the differences between the responses to coherent versus incoherent motion. To obtain a single value that would quantify these asymmetries, the modules of the Fourier coefficients at the first and third harmonics were pooled together using their root-mean-square value (see e.g. [Cottereau et al., 2014]). This operation is equivalent to summing the powers of the individual harmonics and then taking the square root. To take into account the difference of noise levels between the recordings from each of our subjects, we then computed the signal-to-noise ratio (SNR) at odd harmonics by dividing this pooled value by the root mean square of the associated noise magnitudes. Noise magnitude for the first harmonic was given by the average of the moduli at the two neighboring frequencies (i.e 1*F* − *δF* and 1*F* + *δF*, where +*δF*=1Hz is the frequency resolution of our Fourier analysis. Similarly, noise magnitude at the third harmonic was given by the average of the moduli at (3*F* − *δF*)and (3F + *δ*F). For a given subject and a given frequency, the noise magnitude was averaged across all the conditions belonging to the same recording session. The SNRs at even harmonics (that reflect the symmetries in the neural responses to our stimuli) were obtained by repeating the same computation for the second (2F) and fourth (4F) components of the steady-state frequency.

**Figure 2:**
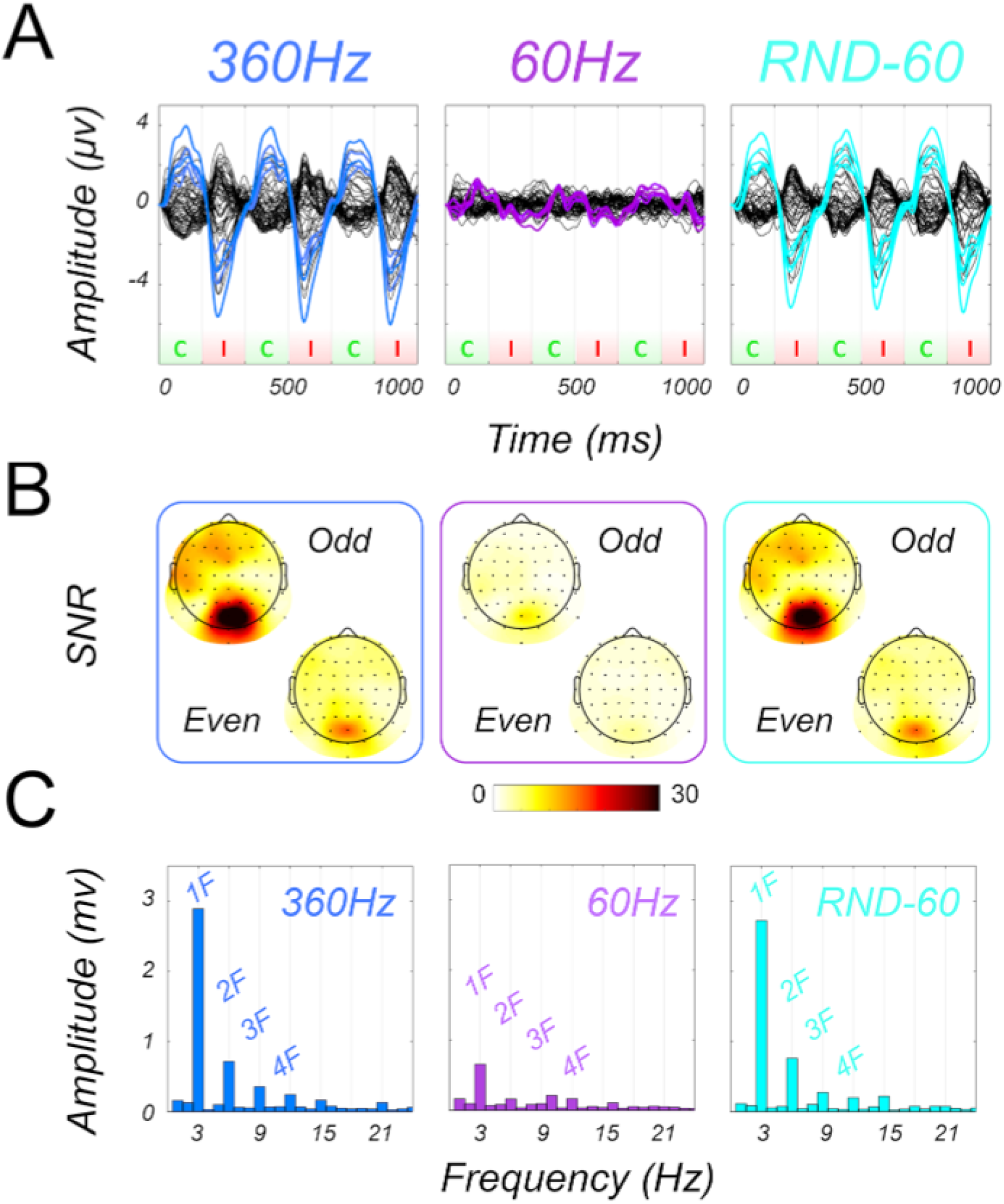
Results of the first experiment in one typical subject. Responses for the 360Hz, 60Hz and RND-60 conditions are respectively shown in the left (in blue), middle (in purple) and right (in cyan) panels. A) Butterfly plot of the responses during one second (i.e. 3 cycles of the 3Hz global motion pattern). The red and green shaded boxes give the alternation between incoherent and coherent motions. The coloured time-courses show the responses in occipital electrodes that were used to compute the power spectra in panel C (see more details in the text). B) Topographic maps of the pooled SNRs at the odd (top) and even (bottom) harmonics. C) Power spectra estimated from the occipital electrodes.

**Figure 3:**
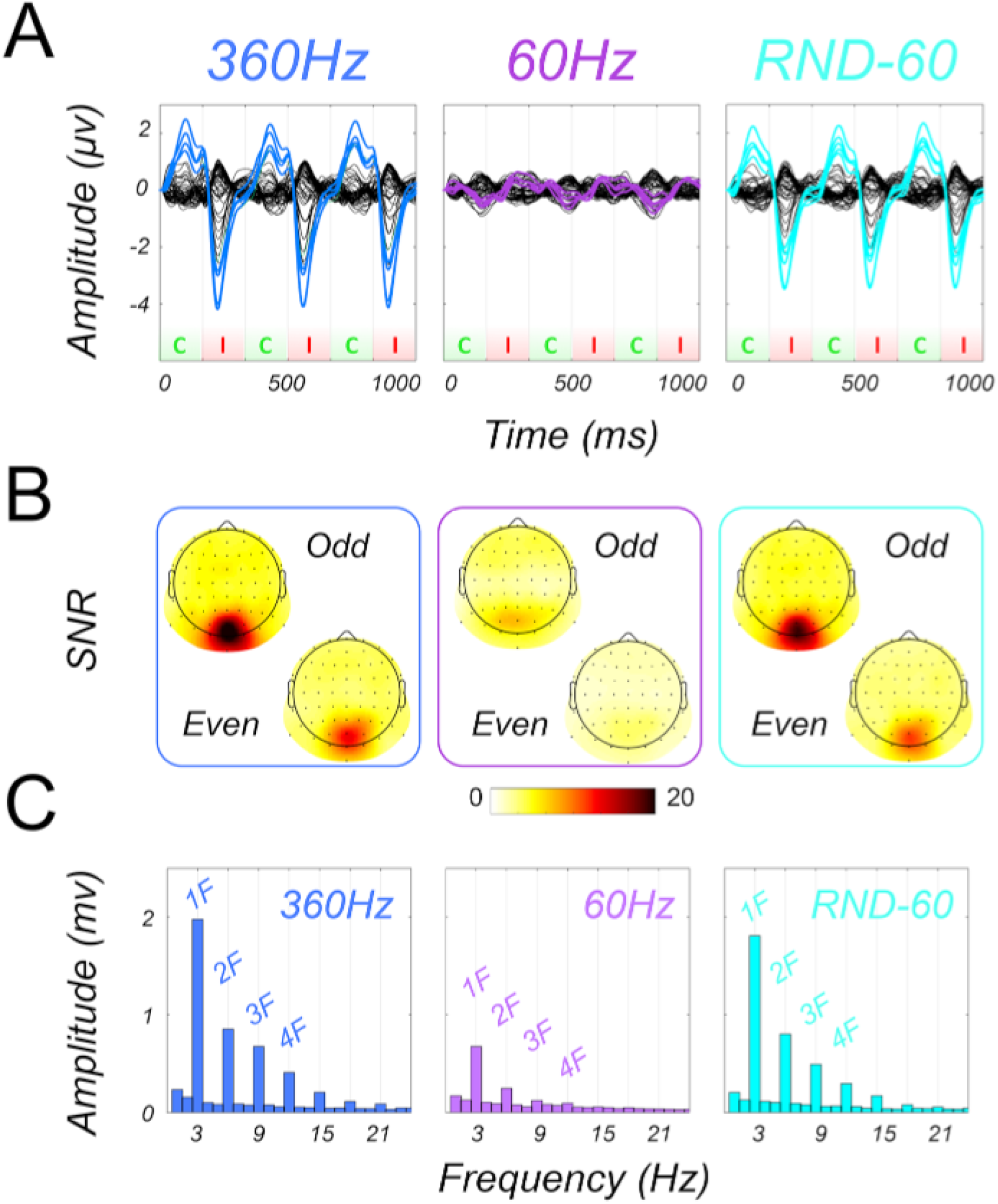
Group-level results of the first experiment (n = 14). Responses for the 360Hz, 60Hz and RND-60 conditions are respectively shown in the left (in blue), middle (in purple) and right (in cyan) panels. See figure 2 for the details of the legend.

### 2.6 Statistical analyses

Based on pilot recordings performed on a few subjects using our 360Hz condition and its 60Hz control, we determined that EEG responses (i.e. the pooled SNRs at odd and even harmonics) to these conditions were stronger in occipital electrodes. Our analyses were therefore performed on the average Fourier coefficients (and consequently pooled SNRs) across these occipital electrodes. Note that because this pilot dataset was not included in the present analysis, our selection of electrodes is therefore independent. This method avoids the double dipping that arises when the same data are used both for identifying electrodes and for measuring activity within them. For a careful comparison between our main condition (360Hz) and the 2 control conditions, we performed a bootstrap analysis. For each subject, we first computed the difference between the pooled SNRs across occipital electrodes in our main versus in our control conditions. The significance of this difference across subjects was evaluated from permutation samples. For each permutation sample, the values corresponding to our two conditions were randomly intermixed before to compute the difference and to average across subjects. We repeated this process for a big number of permutations (2*^N^*, where N is number of subjects). Histogram of all permutation samples was a zero centered distribution whose 95% upper limit was compared to the real difference of pooled SNR between our conditions. We performed this analysis in the two experiments to determine whether the responses in our main condition (360Hz) were stronger than those in our control conditions (60Hz and RND-60 for the first experiment and 120Hz and RND-120 for the second experiment).

## 3 Results

In this study, we were interested in the EEG responses elicited by 3 conditions that contained the same global motion and differed in their local spatio-temporal properties. Figure 2 shows the responses obtained in one typical subject for the first experiment where the control conditions were based on a refresh rate of 60Hz (average responses across subjects are shown in figure 3 for the first experiment and figure 5 for the second experiment).

**Figure 4:**
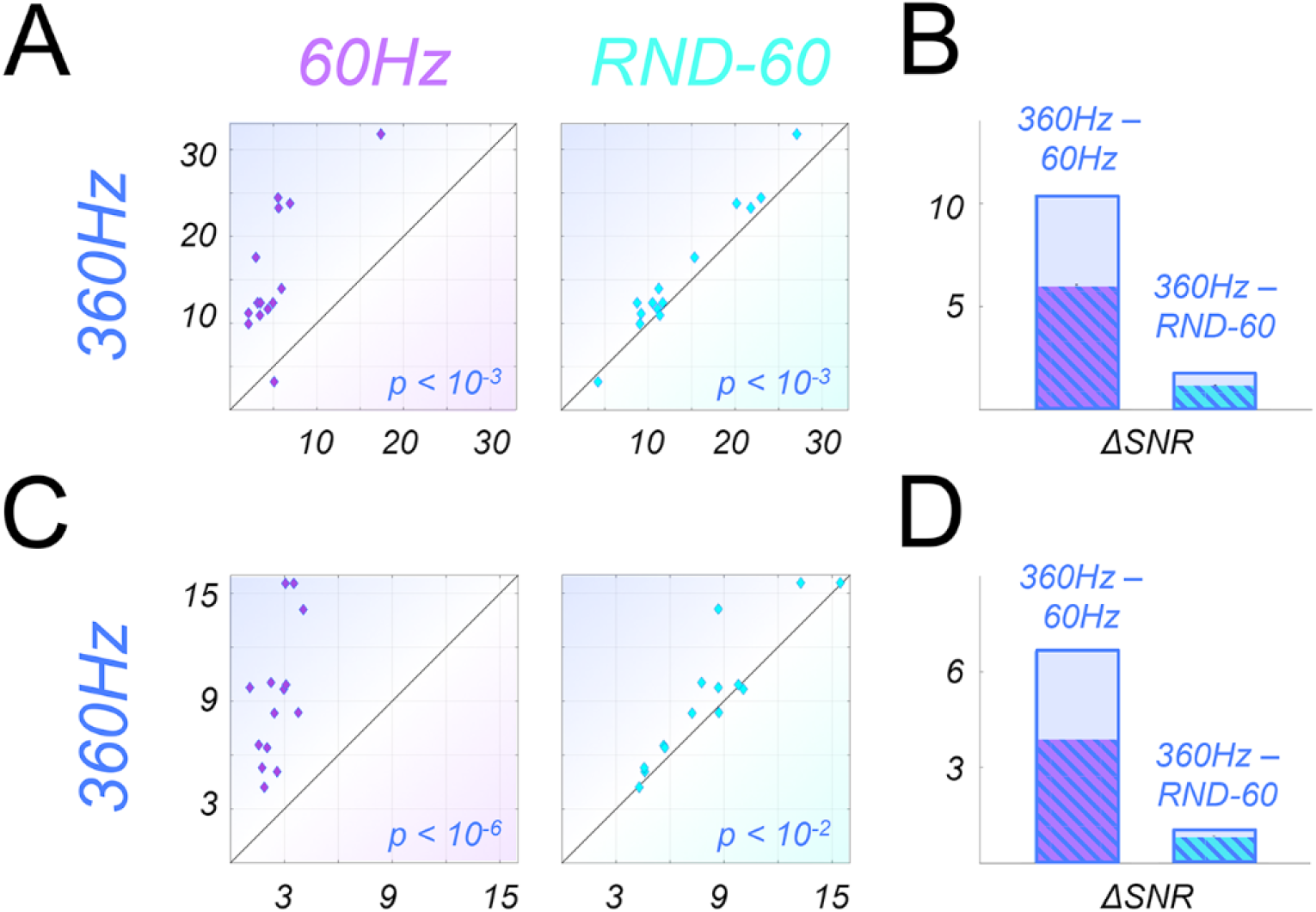
Pooled SNRs in the 360Hz condition as a function of the pooled SNRs in the 60Hz (left panels) and RND-60 (middle panels) conditions. Each diamond corresponds to values measured in one subject. The right panels give the average difference between the 360Hz versus the 60Hz condition (leftward bars) and the 360Hz versus the RND-60 (rightward bars) condition. Hatched areas correspond to the upper limit of the 95% confidence interval of this difference estimated from bootstrap analysis (see the details in the text). The p-values associated with our bootstrap analysis are provided in the lower-right parts of the left and middle panels. A & B) Pooled SNR at odd harmonics. C & D) Pooled SNR at even harmonics.

**Figure 5:**
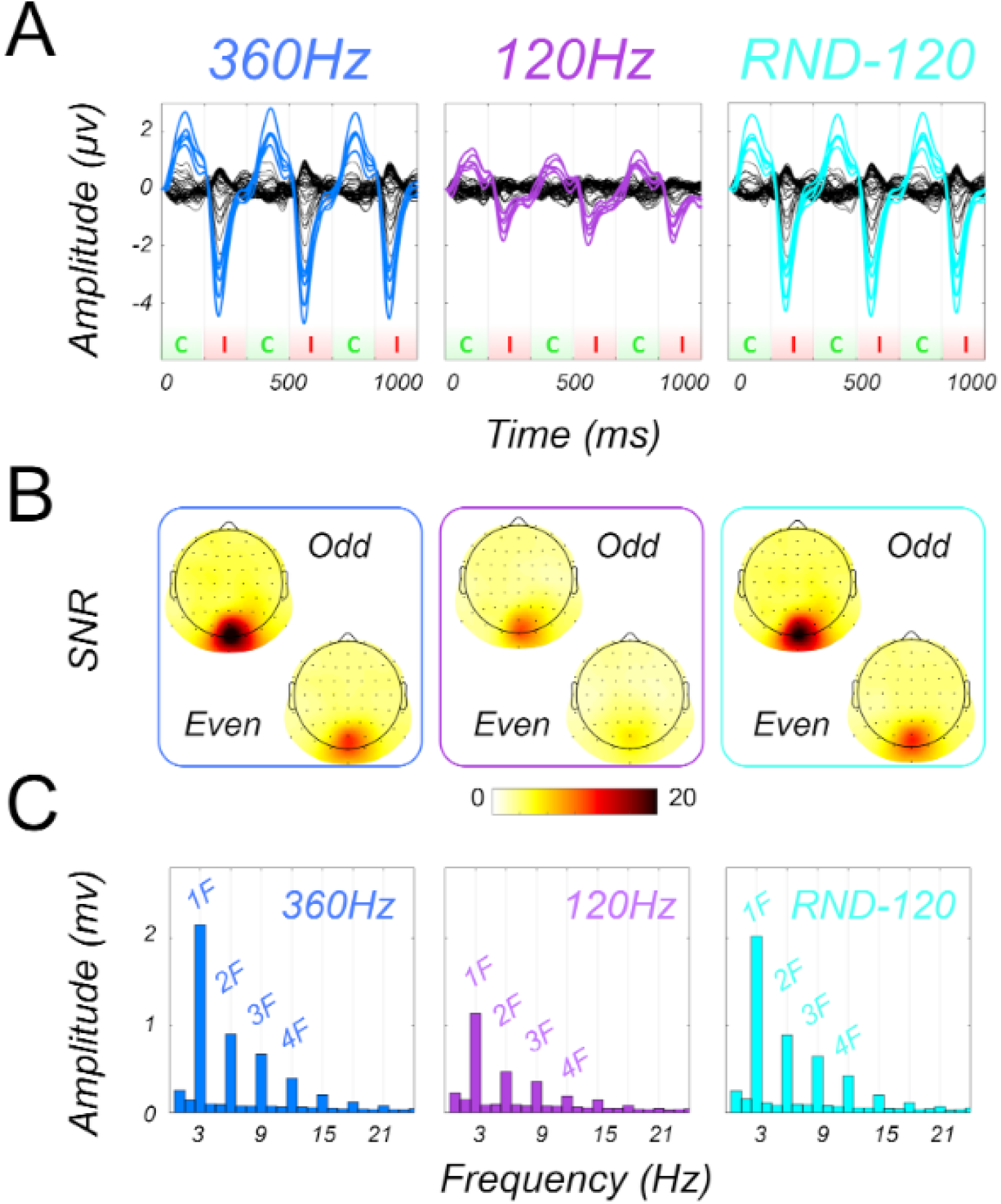
Group-level results of the second experiment (n = 18). Responses for the 360Hz, 120Hz and RND-120 conditions are respectively shown in the left (in blue), middle (in purple) and right (in cyan) panels. See figure 2 for the details of the legend.

The first row (A) shows the butterfly plots (the average data for each electrode) corresponding to 1s of stimulation. Responses corresponding to our 360Hz, 60Hz and RND-60 conditions are respectively shown on the left (in blue), on the middle (in purple) and on the right (in cyan) panels. As in every SSVEP experiment (see e.g. [Norcia et al., 2015]), these responses are frequency encoded. Within one-second time-windows, there are exactly three cycles that correspond to our global motion alternation frequency (i.e. the 3Hz alternation between the coherent and incoherent motion states, see the Materials and methods section). The second rows (B) shows the topographic maps corresponding to the pooled SNRs at the odd (top) and even (bottom) harmonics of the stimulation frequency (see the materials and methods). All three conditions elicit strong activations within occipital electrodes. Finally, the last row (C) shows the power spectra estimated for our three conditions in occipital electrodes. As specified in the materials and methods, these electrodes were selected from pilot recordings performed with the 2 of our stimuli but which nonetheless constitute an independent dataset. Here as well, the frequency content of the responses is expected with stronger responses at the stimulation frequency (3Hz) and its harmonics. Interestingly, if the responses corresponding to our three conditions show similar patterns in this subject, the amplitudes of these responses are not as strong in the 60Hz condition as in the two others (360Hz and RND-60). Responses to the 360Hz are also stronger (although at a lesser degree) that those to the RND-60 condition. This pattern is actually not specific to this particular subject and can also be observed in the group-level data that are shown on figure 3.

In order to characterize whether those differences are significant, we show on figure 4 the pooled SNRs obtained in each subject for the 360Hz condition as a function of those obtained in the same subjects for the 60Hz condition (left panels) and for the RND-60 condition (middle panels).

The two rows correspond to odd (upper row) and even (lower row) pooled SNRs. For all these comparisons, most of the data points lie above the first diagonal (y=x), which means that the responses obtained for the 360Hz condition are stronger than for the other conditions. These observations are confirmed by bootstrap analyses (see the Materials and Methods section) that are presented on the right panels. These panels show the difference between the pooled SNRs for the 360Hz vs 60Hz (leftward bars) and for the 360Hz vs RND-60 (rightward bars) conditions. Here as well, the upper and lower rows correspond to pooled SNR at odd (upper and even (lower) harmonics. The hatched areas provide the upper limit of the 95% confidence interval estimated for those differences. Therefore, the pooled SNRs at both the odd and even harmonics are stronger for the 360Hz condition than for the two control conditions (60Hz and RND-60). The p-values associated with our bootstrap analysis confirmed that these differences were significant (*p* = 1.22 × 10^−4^ and *p* < 10^−6^ for the comparison between the 360Hz condition and its 60Hz control at odd and even harmonics, *p* = 9.6 × 10^−4^ and *p* = 4 × 10^−3^ for the comparison between the 360Hz condition and its RND-60 control at odd and even harmonics). These results demonstrate that brain responses to motion patterns are influenced by the fine-scale spatio-temporal properties of the stimuli.

To determine whether these properties are also observed for control conditions generated from a higher refresh rate, we ran another experiment (n = 18) based on a 120Hz refresh rate (see the Materials and Methods section). Average responses across subjects are shown in figure 5.

Note that the group-level responses to our 360Hz conditions (left panels) are very similar to those observed for the same condition in the first experiment. It shows that our recordings and analyzes are very reproducible. Here again, the responses to the two control conditions (120Hz in the middle panels and RND-120 in the right panels) and to the main condition have similar patterns. Response amplitude are however stronger for the 360Hz condition than for the 120Hz condition. To a lesser degree, responses for the 360Hz condition are also stronger than for the RND-120 condition. These observations were confirmed by our statistical analyses that are shown in figure 6.

**Figure 6:**
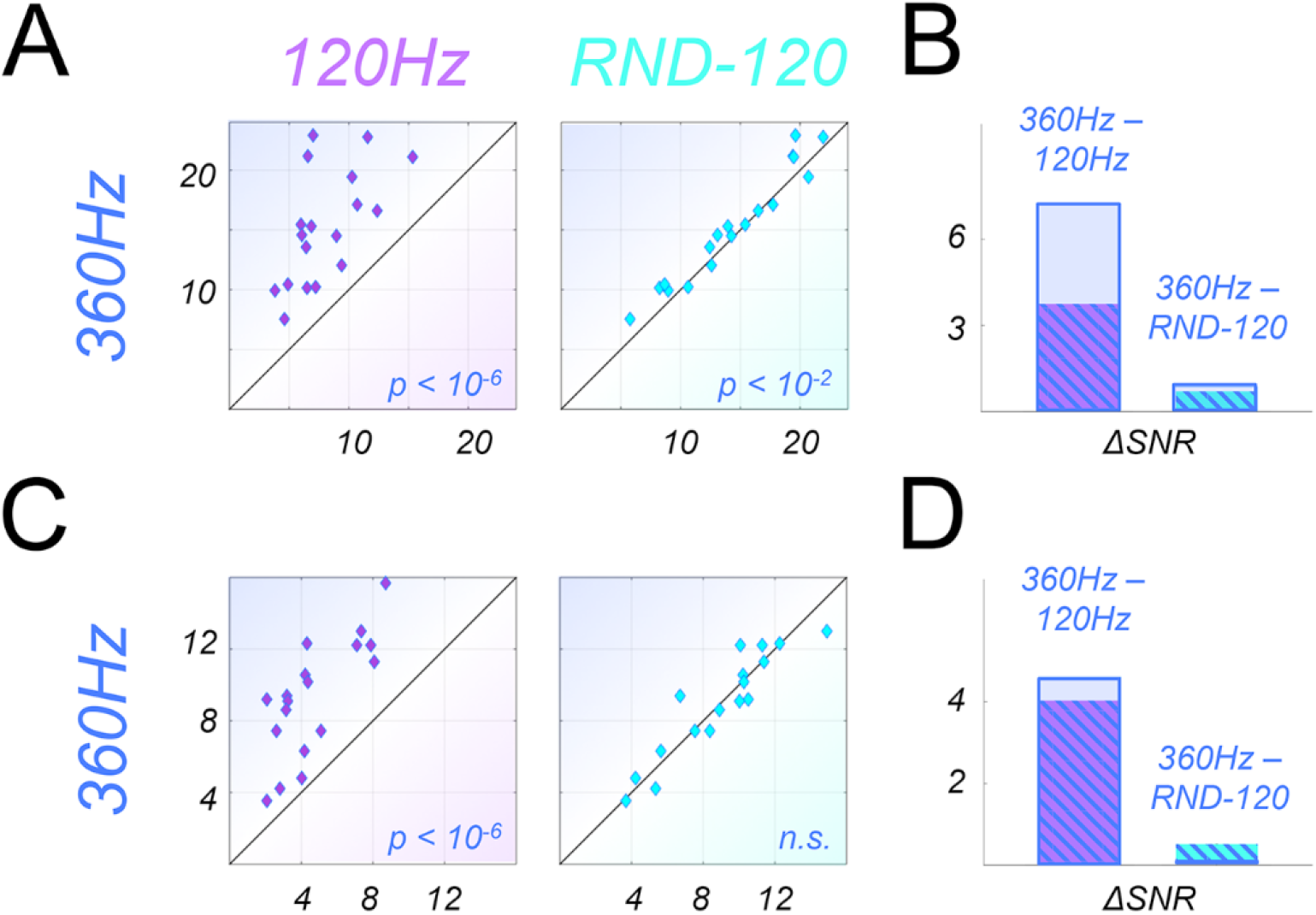
Pooled SNRs in the 360Hz condition as a function of the pooled SNRs in the 120Hz (left panels) and RND-120 (middle panels) conditions (experiment 2). Each dots corresponds to one subject. The right panels give the average difference between the 360Hz versus the 120Hz condition (leftward bars) and the 360Hz versus the RND-120 (rightward bars) condition. Hatched areas correspond to the upper limit of the 95% confidence interval of this difference estimated from bootstrap analysis (see the details in the text). The p-values associated with our bootstrap analysis are provided in the lower-right parts of the left and middle panels. A & B) Pooled SNR at odd harmonics. C & D) Pooled SNR at even harmonics.

The pooled SNR at odd harmonics was significantly stronger for the 360Hz condition than for the two control conditions (*p* < 10^−6^ and *p* = 3 × 10^−3^ respectively). The pooled SNR at even harmonics was stronger for the 360Hz condition than for the 120Hz condition (*p* < 10^−6^). We did not observe significant differences between the pooled SNR at even harmonics for the 360Hz condition versus for the 120Hz condition (*p* = 0.54). If the differences reported here are not as marked as in the first experiment, they are however significant and confirmed that the fine-scale spatio-temporal structure of the stimuli (even based on a 120Hz refresh rate) influence the brain responses to global motion.

## 4 Discussion

The aim of this study was to test whether the fine-scale spatio-temporal structure of motion stimuli influences brain responses recorded with EEG. In a first experiment (n = 14), we characterize the steady-state visual evoked potentials (SSVEPs) to 3 conditions that shared the same global motion pattern (a 3Hz alternation between coherent and incoherent motion) but that differed in their local properties. Our main condition differed from our 60Hz control in its refresh rate and from our RND-60 control in its spatio-temporal profile within a 60Hz frame period. Both these two aspects had an influence on the EEG response amplitude as the pooled SNRs at odd and even harmonics were significantly stronger for the main condition (see figure 2 for the data recorded in one typical subject and figures 3 and 4 for group-level data and analyses). A second experiment (n = 18) where the main condition remained identical but the two controls were based on a refresh rate of 120Hz showed that the local spatio-temporal properties always significantly influenced the SSVEP responses in this case, with stronger responses for the main condition observed at odd harmonics (see figures 5 and 6).

### 4.1 Cortical origins of the recorded signals

EEG recordings mostly reflect responses from large assemblies of pyramidal neurons in cortex [Baillet et al., 2001]. All our motion conditions (and also the difference between our main condition and its controls) elicited strong responses in occipital electrodes (see figures 2, 3 and 5). Our topographic maps are consistent with activations in early visual cortex, e.g. in areas like V1, V2 and V3 (see [Ales et al., 2010]).

However, this does not rule out that other cortical regions in higher visual cortex were also activated during our experiment. In a previous study, using stimuli that also contained translational motion, we obtained very similar topographic maps and we were able to demonstrate with fMRI-informed source imaging that several regions of the dorsal pathway (including the motion-sensitive complex hMT+) were also activated in addition to early visual cortex [Cottereau et al., 2014].

### 4.2 Implication for screen refresh rates

Because current CRT display monitors have refresh rates well above the flicker fusion limit (24fps), it is sometimes believed that they are sufficient to elicit a stable perception of dynamic scenes. The significant differences that we found between the responses evoked by our main condition and those elicited by our 60Hz (experiment 1) or 120Hz (experiment 2) controls actually demonstrate that classical monitors are not optimal in their abilities to activate the visual system as strongly as it is for stimuli with much higher refresh rate that are closer to the real (i.e. continuous) motion stimuli we experience in our everyday life. Pilot data with the same control as here but based on a frame rate at 240Hz (see figure 7) suggest that even state-of-art monitor are not enough to generate strong EEG activations.

**Figure 7:**
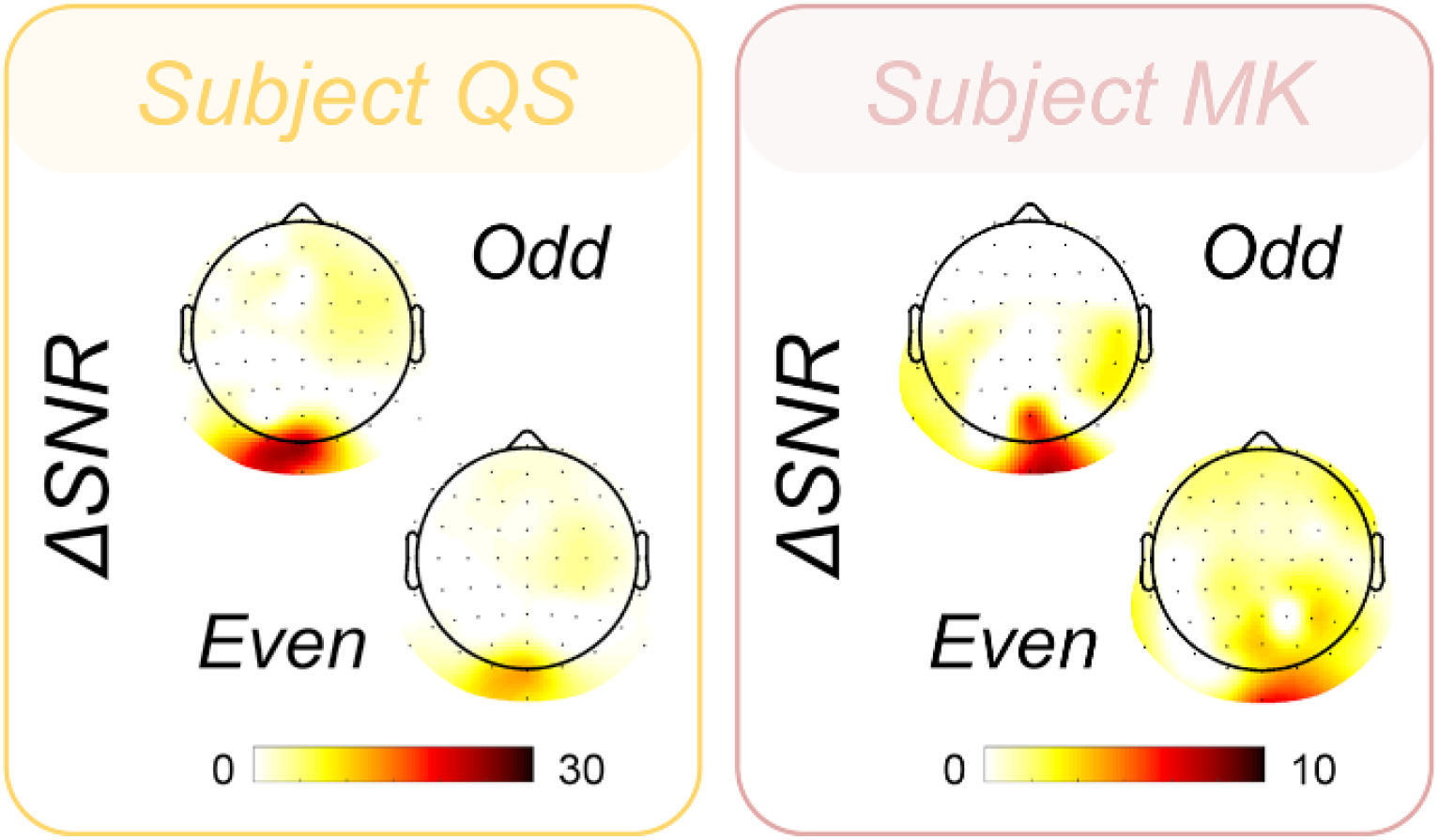
Difference of pooled SNRs between responses to our 360Hz condition and a control at 240Hz in two subjects (subjects QS and MK). Topoplots are shown for both odd and even pooled SNR difference.

Even though our subjects were only involved in a passive fixation, the modification of neural responses that are reported here could very well also influence visual perception. Indeed, using subjective psychophysical reports, [Kuroki et al., 2007] showed that high frames rate improved motion image quality. [Claypool et al., 2006] showed that frame rates impacted performances of users playing video games. From point of view of perceptual information loss [Nasiri and Wang, 2017] have reported that reduced frame rate leads to degraded video quality. More recently, we showed that these differences in refresh rate also influenced different aspects of perception such as speed discrimination or reading abilities [Kime et al., 2016]. The use of monitors with higher frame rates (≥ 240*Hz*) therefore appears as very important to elicit stronger cortical responses and visual perception.

### 4.3 Implication for models of the retino-geniculate pathway

One possible issue with the 60Hz (experiment 1) and 120Hz (experiment 2) control conditions is that the global amount of pixels that were stimulated along the motion pathway is smaller than in the main condition (see figure 1). As a consequence, the differences in signal amplitude found in the experiments might mainly be driven by the numbers of photoreceptors that were activated (note nonetheless that this point does not affect the conclusion of the previous section about the importance of using high refresh rate in displays). This hypothesis is ruled out by the significant differences that we found between our main conditions and the RND-60 control condition. In this case, the refresh rates and spatial positions of the dots within a frame were exactly identical and only the temporal orders differed. However, we still found significantly stronger responses in our main condition than in our controls. It demonstrates that the precise temporal structure within a 60Hz frame (i.e. within a 16.7ms time window) and within a 120Hz frame (i.e. within a 8.3ms time window) influences EEG responses. Because EEG responses mostly reflect responses from early visual cortex (and possibly higher-level cortical areas, see above), it implies that the temporal precision of the neural signals transmitted along the retino-geniculate pathway is below 10ms. This result goes against models that posit that the retino-geniculate pathway actually low-pass filters the visual information so that the precise temporal information is lost when it reaches early visual cortex. For example, for high-speed and high-contrast motion, [Chichilnisky and Kalmar, 2003] estimated a filter time constant of approximately 10ms. Our results are in line with previous studies on animal models that reported responses with temporal resolution near the millisecond in different part of the visual pathway. Recording from RGCs on rabbits and salamanders in response to random flickers demonstrated transient and sparse spike bursts, with highly reproducible timings [Berry et al., 1997]. Millisecond precision has been observed on spiking responses of cat LGN by presenting repeated white noise patterns, with a very low inter-trial variability [Reinagel and Reid, 2002]. This very precise spiking activity was robust across cells of the same type. Importantly, if it was shown that the relative temporal precision of LGN responses can vary depending on the content of the stimulus [Butts et al., 2007], in vivo recordings in cat LGN showed that responses to drifting sinusoidal gratings had temporal precision around 5ms for high-contrast values [Reich et al., 1997], similar to those used in the present study.

In addition to the presence of responses with high-temporal resolution along the retino-geniculate pathway, our data also suggest that neurons in early visual cortex have spatiotemporal receptive fields that favor continuous motion, even within very small temporal windows (i.e. smaller than 8.3ms) because EEG responses were stronger in our main condition.

